# Experimental Study of Social Signaling through Delayed Plumage Maturation in a Colony-Nesting Seabird

**DOI:** 10.1101/2025.09.05.674431

**Authors:** Molly M. Hill, Sarah L. Dobney, Liliane K. Fanburg, Daniel J. Mennill, Liam U. Taylor

## Abstract

Delayed development is a widespread evolutionary strategy that can reduce competition among highly social animals. Many seabirds exhibit delayed plumage maturation, in which young birds spend years in visually distinct predefinitive plumages before attaining the definitive plumage of adults. Previous work hypothesized that predefinitive plumages may function to reduce aggression towards young, predefinitive seabirds at breeding colonies, an idea known from other lineages of birds as the status signaling hypothesis. We tested this hypothesis with visual stimulus experiments at a breeding colony of American Herring Gulls (*Larus smithsonianus*). We presented painted models of four different plumage classes (first-cycle predefinitive gull plumage, third-cycle predefinitive gull plumage, definitive gull plumage, and a Canada Goose as a control) and measured the aggressive responses of breeding adults at their nests. Breeding gulls responded with significantly less frequent, lower, and slower aggression toward the first-cycle and control plumages compared to the definitive plumage. There were no significant differences in response towards the third-cycle plumage. These results offer support for a status signaling hypothesis, indicating that substantial, mottled brown plumage—as worn by first-cycle gulls—reduces aggression from breeding adults in colonies or foraging flocks. Future research can investigate how immature seabirds—including third-cycle gulls—may combine plumage, posture, and behavior to shift the dynamics of social behavior at breeding colonies.

## Introduction

One evolutionary consequence of sociality is a delayed developmental form that reduces competition. For example, orangutans reduce the threat of conflict by suppressing the development of male face flanges (Kralick et al. 2023) while some highly social wrasses delay developing male sexual characteristics until potential competitors are absent (Warner and Swearer 1991). In birds, several lineages have evolved delayed plumage maturation, in which individuals spend multiple years in distinctive predefinitive plumages before acquiring definitive plumages with adult coloration (Hawkins et al. 2012). Predefinitive plumages can act as social signals that reduce aggression towards young birds (Lyon and Montgomerie 1986, Hawkins et al. 2012), such as for Long-tailed Manakins (*Chiroxiphia linearis*) at display leks (McDonald 1993) and Lazuli Buntings (*Passerina amoena*) at breeding territories (Muehter et al. 1997). Through behavioral studies of animals’ reactions to different developmental stages, we can better understand the dynamics of animals’ lives in complex social environments.

Here, we investigate the social signaling function of delayed plumage maturation in seabirds at breeding colonies. Prolonged delayed plumage maturation, lasting two years or longer, is present in multiple, independent lineages of seabirds including gulls (Charadriiformes: Laridae), albatrosses (Procellariiformes: Diomedeidae), and gannets (Suliformes: Sulidae). Many species in these lineages have predefinitive plumages that are distinctly darker or drabber than definitive adult plumages (Grant 1982, Tickell 2000, Nelson 1978). The ecological, evolutionary, and behavioral properties of delayed plumage maturation in seabirds are not yet clear.

Behavioral observations of seabirds have raised a functional hypothesis for delayed plumage maturation in these lineages. In many seabird species, immature birds return to nesting colonies while still in a distinctly predefinitive plumage, despite not breeding for multiple additional years (e.g., Pickering 1989, Nelson 1966). Young birds in predefinitive plumages also occur alongside adult birds in foraging flocks away from breeding colonies (Wakefield et al. 2019). Accordingly, a functional hypothesis is that predefinitive plumages are social signals that reduce the risk of conflict for young seabirds when they start visiting colonies (Taylor 2024) or competing in foraging flocks. This idea corresponds to the status signaling hypothesis first presented in relation to songbirds (Lyon & Montgomerie 1986). Alternatively, delayed plumage maturation may be socially neutral (i.e., it may serve no social signaling function) if it is merely the byproduct of delayed hormonal expression in young, non-breeding birds (Schaedler et al. 2021).

We tested the status signaling hypothesis with a model presentation experiment at a breeding colony of American Herring Gulls (*Larus smithsonianus*). We simulated conspecific birds in different plumage stages near the nests of breeding gulls, using plastic models painted to match different predefinitive and definitive gull plumages, as well as a heterospecific control stimulus. We measured the aggressive responses of breeding gulls to presentations of (1) a young, first-cycle predefinitive gull plumage, (2) an adolescent, third-cycle predefinitive gull plumage, (3) a definitive gull plumage, and (4) a Canada Goose as a control. We predicted that nesting gulls would respond most aggressively to the definitive models and less aggressively to the first-cycle, third-cycle, and control models based on the status signaling hypothesis. Alternatively, if predefinitive plumages are socially neutral, we predicted that breeding gulls would respond with equal aggression levels to all gull plumage models. This experiment provides an opportunity to investigate how predefinitive plumages may function as social signals to mediate aggression at seabird colonies.

## Methods

### Study species and site

American Herring Gulls (*Larus smithsonianus*) are seabirds that breed in coastal, urban, and island colonies across much of northern North America. The species exhibits delayed reproduction and delayed plumage maturation, taking approximately four years to begin breeding (as described for the closely-related European Herring Gull *Larus argentatus*; Chabrzyk and Coulson 1976) in definitive adult plumage (Grant 1982).

The definitive plumage in the American Herring Gull has white body feathers with pale gray wings, black wing tips with white “mirror” patches, a bright yellow bill, and a red gonydeal spot near the tip of the bill (Fig. 1A). Before reaching this definitive plumage, young birds spend approximately one year in each of three predefinitive plumage stages: (1) a mottled brown plumage in the first molt cycle, ∼12 months old (Fig. 1B); (2) a highly variable, mottled brown plumage with a gray saddle in the second molt cycle, ∼24 months old; and (3) a near-definitive plumage featuring gray wings with brown coverts, dark brown primaries, a dark tail band, and a yellow bill with merged black and red spots near the tip in the third molt cycle, ∼36 months old (Grant 1982, Nisbet et al. 2017, Taylor 2024; Fig. 1C).

**Figure 1.**
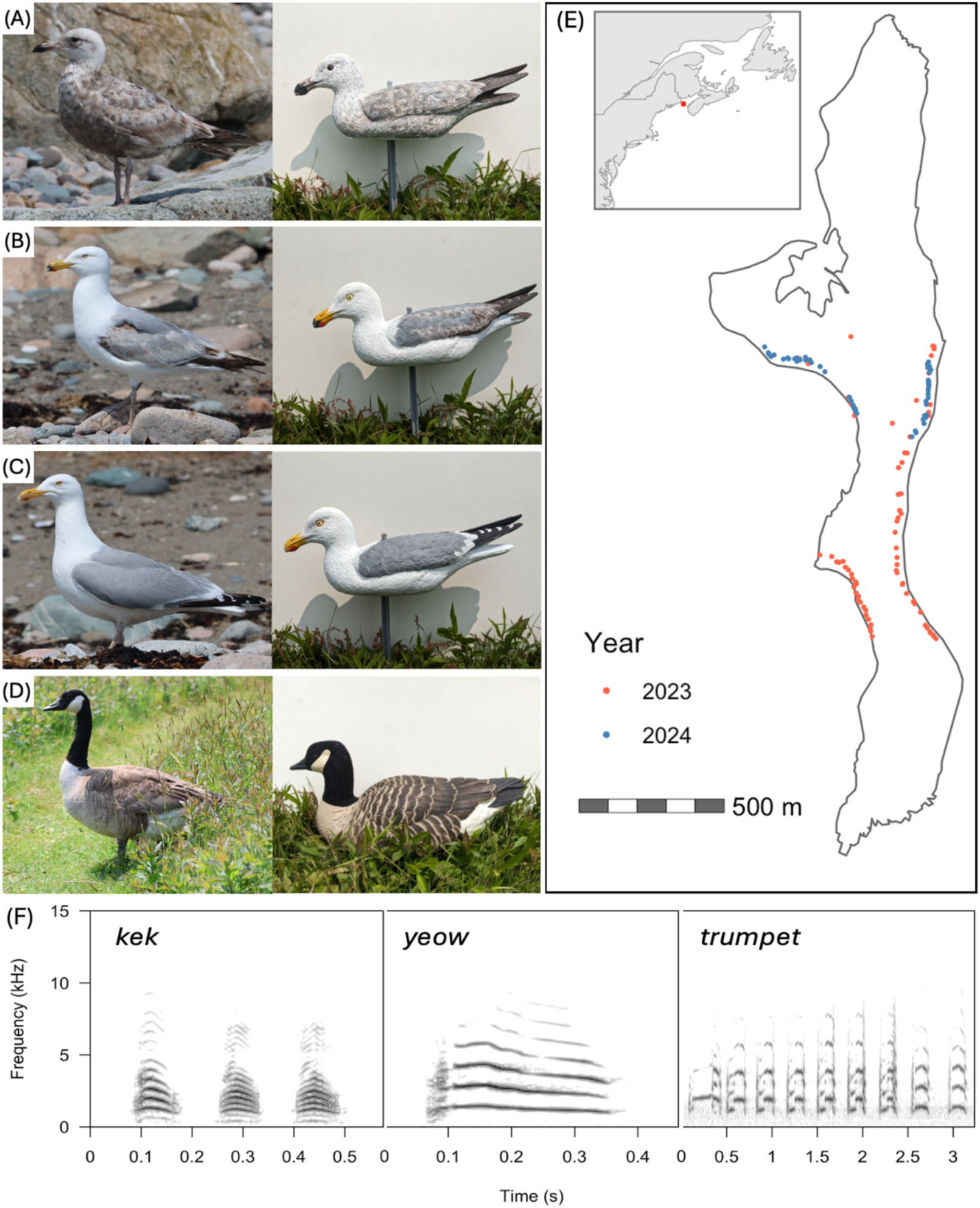
Experimental stimuli and study site for an investigation of plumage signals in American Herring Gulls (*Larus smithsonianus*). Photographs of (A) first-cycle gull plumage (∼12 months old); (B) third-cycle gull plumage (∼36 months old); (C) definitive gull plumage (46+ months old); and (D) Canada Goose (control), including live birds photographed on Kent Island (left) and photographs of one model of each type. (E) A map of the American Herring Gull breeding colony on Kent Island, New Brunswick, Canada, showing the location of nests used in experimental trials. (F) Sound spectrograms showing an example of each of the calls used in the acoustic stimuli.

We conducted our study at the American Herring Gull breeding colony on Kent Island in the Bay of Fundy, New Brunswick, Canada (latitude 44.583, longitude –66.756; Fig. 1E). This 80-hectare island consists of a large grass field along with patches of spruce and deciduous forest. Roughly 6,000 gulls breed on the island each year (Bennett et al. 2017). Gulls begin arriving to Kent Island from wintering grounds in the eastern and southeastern United States in the early spring (Gross 1940, Anderson et al. 2019). They establish nesting territories by early May. The incubation period at the site generally runs from mid-May through mid-June, during which time birds are highly territorial and defend the areas around their nests (Tinbergen 1953, Nisbet et al. 2017). Both parents assist with incubation, with partners alternating between attending the nest and leaving the territory to forage (Shlepr et al. 2021). Most gulls on Kent Island are breeding birds in definitive plumage, but a small number of first– and second-cycle birds (<1% of the total population) along with a larger number of third-cycle birds (∼4% of the total population) are regularly found within the colony as prebreeders (Taylor 2024).

### Primary visual stimuli

Our experiment used four visual stimuli as different plumage treatments. Three were plastic gull models (model: SG 1100; Sport Plast Decoy Co., Medelana, Italy). The models were hand-painted with acrylic paint to match different American Herring Gull plumage stages: first-cycle, third-cycle, and definitive (Fig. 1A–C). The control stimulus was a plastic Canada Goose (*Branta canadensis*; model: AVX-LGCF-4PK; Avian-X Decoys,

Irving, TX, USA), which is a common breeding species on Kent Island that does not regularly exhibit aggressive interactions with gulls (Fig. 1D). There were three exemplars of each gull model and two exemplars of the goose model, for a total of 11 visual stimuli (Fig. S1). To minimize the effects of pseudoreplication, exemplars were randomized across trials.

### Secondary acoustic stimuli

We used acoustic stimuli to help draw attention to the visual stimuli. Acoustic stimuli consisted of three call types presented in an order corresponding to increasing levels of agitation: *kek* indicating mild alarm, *yeow* indicating moderate alarm, and *trumpet* indicating territoriality and social aggression (Tinbergen 1953; Nisbet et al. 2017). Each acoustic stimulus had the following structure: 20.0 s of silence; three repetitions of a cluster of three *kek* calls, each separated by 3.0 s of silence; 10.0 s of silence; three repetitions of a *yeow* call, each separated by 3.0 s of silence; 10.0 s of silence; three repetitions of a *trumpet* call, each separated by 3.0 s of silence (Fig. 1F). We created these stimuli from recordings we collected on Kent Island in August 2022. Recordings were collected with an ME62/K6 microphone mounted in a Telinga parabola connected to a Marantz PMD-661 digital recorder (44.1 kHz sampling rate; 16-bit accuracy; WAV format). We selected high signal-to-noise ratio recordings of different individual birds, recorded in different locations throughout the island, producing *kek*, *yeow*, and *trumpet* calls.

We prepared playback stimuli using Adobe Audition using three exemplars for each of the three call types. *Kek* calls are often uttered in clusters of three notes spaced closely together (Nisbet et al. 2017). For *kek* calls, we ensured that each recording had three notes, spaced approximately 0.1 s apart, for a total duration of approximately 0.5 s. For *yeow* calls, we used individual calls of approximately 0.3 s in duration. For *trumpet* calls, we used individual calls, each approximately 3.0 s in duration and containing 9 notes. We filtered background sounds out of each recording using FFT filters and the LASSO tool in Audition. We then normalized the amplitude of each recording to –1 dB. We created 27 different audio tracks using all possible combinations of the three recordings of each call type. To minimize pseudoreplication, tracks were randomized across trials. We broadcast audio using a camouflage JBL Flip 5 speaker at a consistent volume that we determined to be natural in the field based on aural assessment. Later, using a calibrated Casella CEL-24X sound level meter, we measured the amplitude of each call type, and found *kek* calls were broadcast at 72-76 dB SPL, *yeow* calls were 75-77 dB SPL, and *trumpet* calls were 82-86 dB SPL, measured at 1m distance from the loudspeaker.

### Nest selection

We selected nesting territories for each trial in easily accessible field and beach habitat (Fig. 1E), choosing only nests with a parent in attendance. Most selected nests had complete, three-egg clutches (113 / 120 nests) while the remainder had two eggs. To account for potential variation in parental aggression due to differential investment in the developing clutch (Pierotti and Annett 1994), we selected only nests that were predicted to be within one week of hatching based on egg floatation (Schreiber 1970). Although some eggs began hatching across multiple trials at a single nest, no chicks left the nest area across the days when we presented the stimuli to the nesting gulls.

### Experimental procedure

We tested the aggressive response toward gull plumage signals by presenting models near the subjects’ nests, using the four plumage treatments at each nest in a randomized order. Each trial at a nest was separated by 24–36 hr to minimize disturbance and habituation effects. With 120 nests and four plumage treatments per nest, the total number of trials was 480.

For each trial, one observer approached the nest with the visual stimulus and the loudspeaker. Models were set upright on a freely spinning, plastic spike ∼0.75 m from the nest and the loudspeaker was placed underneath (Fig. S2). That observer started the playback stimulus and began briskly walking away from the nest, at which point a second observer started a timer. These two observers helped categorize the behavioral responses of parents along with the time at which each behavioral response first occurred. Nest defense behaviors can vary when both parents are present at the nest (Shah et al. 2025), so we also recorded the number of parents present at each trial (one, or two, or zero if adults left the area at the start of the trial). Because many trials featured only one parent, we did not analyze the sex of these unmarked birds using visual comparisons (the sexes are monochromatic, and although males are larger than females, an observer can only make such a differentiation if both parents are near one another). There were no cases where parents from neighboring nests directly interfered with the stimuli. Each trial lasted 2 min, or until a bird made physical contact with the visual stimulus. To prevent destruction of the model, we stopped a trial early and removed the stimulus when a bird made physical contact.

### Behavioral scoring

We categorized behavioral responses into five aggression levels (Table 1). Level 0 (Relaxed) was reserved for gulls that exhibited relaxed behaviors such as preening or incubating. Level 1 (Alarm) included agitation or alarm responses (*e.g.*, *kek* calls, *yeow* calls, rapid pacing, short fluttering flights). Level 2 (Social Aggression) included territorial or social conflict displays such as *mew* calls, *trumpet* calls, and grass-pulling displays directed at the model (Tinbergen 1953). Level 3 (Physical Threat) included lunging or swooping at the model without making direct contact. Level 4 (Physical Attacks) included direct contact with the model.

**Table 1.** Behavioral aggression levels used for analysis.

| Level | Category | Description |
| --- | --- | --- |
| 0 | Relaxed | Incubation; preening; standing; regurgitating; eating |
| 1 | Alarm | Pacing; rapidly approaching stimulus; fluttering flight; *kek* or *yeow* vocalizations |
| 2 | Social aggression | Choking displays; grass-pulling displays; *trumpet* or *mew* vocalizations |
| 3 | Physical threat | Lunging or swooping at stimulus |
| 4 | Physical attack | Contact with stimulus |

Each trial was assigned a single aggression score, corresponding to the maximum aggression level observed during that trial. We excluded behavioral responses that were clearly directed towards the observers or towards neighboring birds, retaining other responses for that trial. We excluded trials where birds flushed at the start of the trial and never returned. After these filters, there was a total of 442 trials retained for analysis: 109 trials for the goose control treatment, 110 trials for first-cycle gull plumage, 113 trials for the third-cycle gull plumage, and 110 trials for the definitive gull plumage.

### Data analysis

All analyses were conducted using the *tidyverse* packages in R version 4.4.1(Wickham et al. 2019; R Core Team 2024). First, we used a binomial logistic model to test whether the rate of non-aggressive responses (maximum aggression levels 0–1) versus aggressive responses (maximum aggression levels 2–4) differed by plumage treatment. The model included treatment order (1, 2, 3, or 4) and number of parents present as fixed effects, and nest ID as a random effect (Fig. S3). We fit the model using the *glmer* function from the *lme4* package (Bates et al. 2015). The intercept categories were the definitive plumage treatment, treatment order 1, and a single parent in attendance for all models.

To investigate different levels of aggression in more detail, we also fit an ordinal mixed effects model. The ordinal model allowed our five aggression levels to be increasing in relative intensity without assigning them precise quantitative values. As in the previous model, we included treatment order and number of parents as fixed effects along with nest ID as a random effect. We fit the model using the *clmm* function from the *ordinal* package (Christensen 2023).

Finally, we tested for differences in speed of aggressive responses using a linear mixed effects model. The response variable in this model was the minimum time needed to reach the maximum aggression level within each trial. We only included trials where there was an aggressive response (i.e., maximum aggression levels 2–4). In addition to treatment order (fixed effect), number of parents (fixed effect), and nest ID (random effect), this model also included aggression level itself (2, 3, or 4) as a fixed effect. We fit the model using the *lme* function from the *nlme* package (Pinheiro and Bates 2024)

### Ethical Note

All experimental procedures were approved by the Bowdoin Scientific Station, Bowdoin College (Protocol #2023-0008), and Yale University (Protocol #2024-20453). Birds were not captured or handled for this experiment. We designed the experiment to minimize the length of exposure of the animals to signals that could be perceived as aggressive.

## Results

Nesting American Herring Gulls responded with variation in their rate of aggressive response to visual stimuli of different gull plumage stages (Table 2; Fig. 2). Across all plumage treatments, 67% of trials resulted in a non-aggressive response (298 / 442 trials; maximum aggression levels 0–1) while 33% of trials resulted in an aggressive response (144 / 442 trials; maximum aggression levels 2–4). Based on binomial mixed effect model estimates, gulls were 48% less likely to respond aggressively to the first-cycle gull plumage treatment (p < 0.05) and 86% less likely to respond aggressively to the goose control (p < 0.001) when compared to definitive gull plumage (Table 2). We found no evidence differences in the rate of aggression towards the third-cycle plumage treatment, nor were there effects of treatment order or number of parents present during the trial (Table 2).

**Figure 2.**
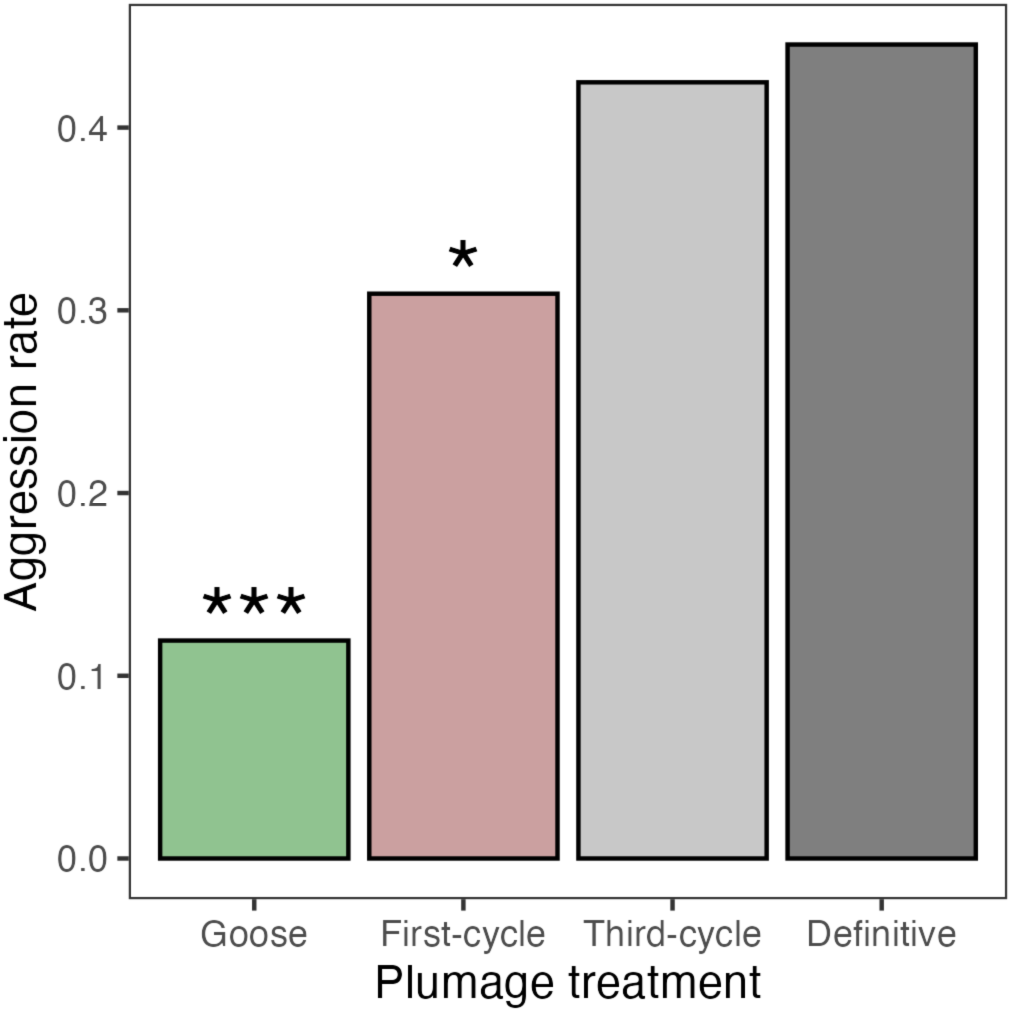
American Herring Gulls were less likely to respond aggressively toward first-cycle gull plumage, and Canada Goose control models, compared to the definitive gull plumage. Aggression rate was calculated as the percentage of trials reaching a maximum aggression level of 2–4 for each treatment. Asterisks indicate p < 0.05 (*) and p < 0.001 (**) compared to the intercept category of definitive plumage, from a binomial mixed effects model.

**Table 2.** Results of binomial mixed effects model for aggressive response rate. Trials were considered non-aggressive if they reached maximum aggression levels 0–1 and aggressive if they reached levels 2–4. Definitive gull plumage treatment is the intercept category.

| **Model term** | **Estimate** | **Odds ratio** | **S.E.** | **z** | **p** |
| --- | --- | --- | --- | --- | --- |
| (Intercept) | -0.35 | 0.70 | 0.30 | -1.17 | 0.24 |
| Plumage – Third-cycle | -0.08 | 0.92 | 0.29 | -0.28 | 0.78 |
| Plumage – First-cycle | -0.66 | 0.52 | 0.30 | -2.20 | **0.03** |
| Plumage – Goose | -1.96 | 0.14 | 0.38 | -5.20 | **<0.001** |
| Trial order – 2 | -0.15 | 0.86 | 0.32 | -0.46 | 0.65 |
| Trial order – 3 | -0.05 | 0.95 | 0.33 | -0.15 | 0.88 |
| Trial order – 4 | 0.46 | 1.58 | 0.32 | 1.45 | 0.15 |
| Num. parents – 2 | 0.15 | 1.16 | 0.32 | 0.47 | 0.64 |

Similarly, the ordinal model revealed lower aggression towards the first-cycle gull plumage, along with the goose control, relative to the definitive gull plumage (Table 3; Fig. 3). In terms of the maximum aggression level, across all trials, 21.0% peaked at no aggression (level 0) while 46.4% peaked at alarm response (level 1), 4.5% at social aggression (level 2), 12.2% at physical threat (level 3), and 15.8% at physical attack (level 4; Fig. S4). We found no evidence for relationships between aggression level and third-cycle gull plumage, treatment order, or number of parents (Table 3, Fig. 3).

**Figure 3.**
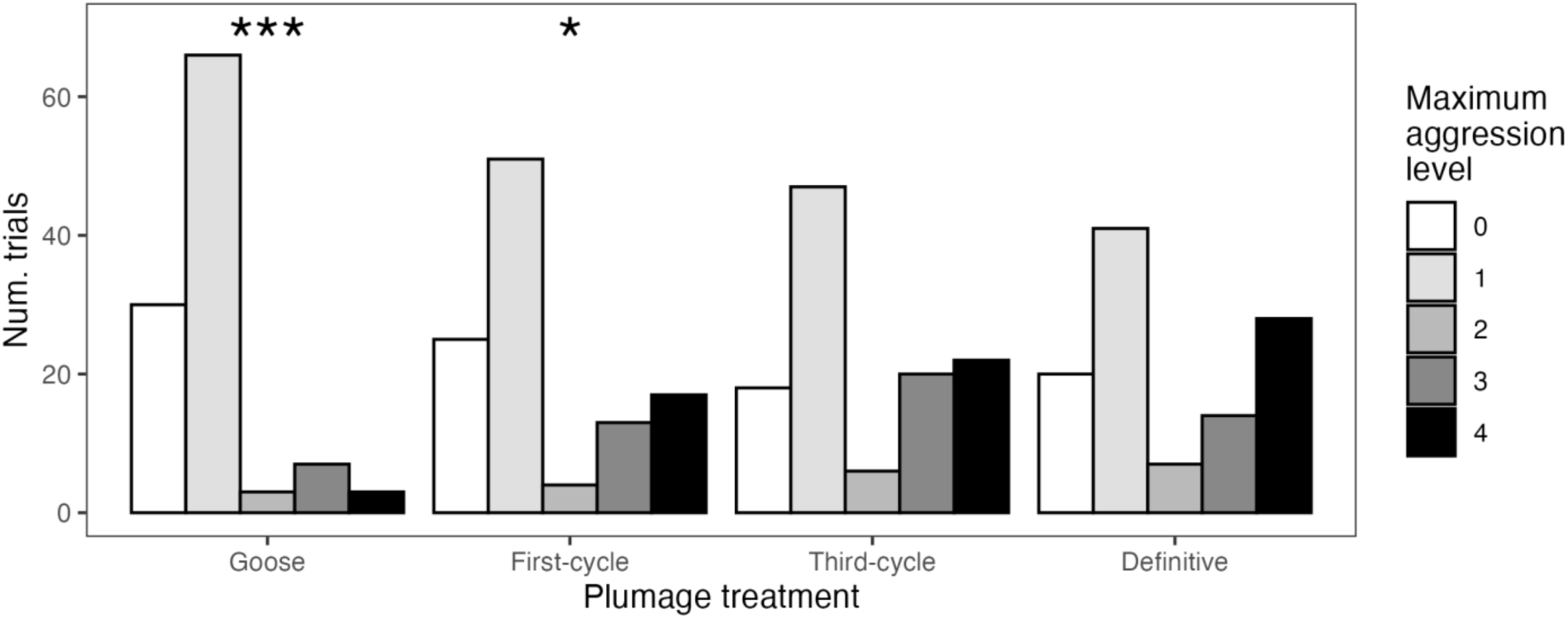
American Herring Gulls responded with lower aggression towards the first-cycle gull plumage, and Canada Goose control, compared to the definitive gull plumage. Vertical lines indicate the mean aggression level for each plumage treatment. Asterisks indicate p < 0.05 (*) and p < 0.001 (***) compared to the intercept category of definitive plumage, from an ordinal mixed effects model.

**Table 3.** Results of ordinal model for maximum aggression level (0, 1, 2, 3, or 4). Definitive gull plumage treatment is the intercept category.

| **Model term** | **Estimate** | **S.E.** | **z** | **p** |
| --- | --- | --- | --- | --- |
| Plumage – Third-cycle | -0.02 | 0.26 | -0.09 | 0.93 |
| Plumage – First-cycle | -0.57 | 0.27 | -2.16 | **0.03** |
| Plumage – Goose | -1.33 | 0.27 | -4.95 | **< 0.001** |
| Trial order – 2 | -0.35 | 0.26 | -1.34 | 0.18 |
| Trial order – 3 | -0.41 | 0.26 | -1.58 | 0.12 |
| Trial order – 4 | 0.03 | 0.26 | 0.13 | 0.89 |
| Num. parents – 2 | 0.43 | 0.26 | 1.65 | 0.10 |

When analyzing aggressive responses only (i.e., response levels 2 and higher), gulls took significantly longer to reach maximum aggression towards the first-cycle gull plumage (mean response time: 43.9 ± 36.1 s) compared to the definitive gull plumage (36.7 ± 27.7 s; Table 4; Fig. 4). We found no evidence for effects on timing from the third-cycle gull plumage (34.9 ± 30.2 s), and a large but statistically insignificant effect from the goose plumage (50.8 ± 37.6 s), although the power of the this comparison is likely limited by the small number of aggressive responses towards the goose.

**Figure 4.**
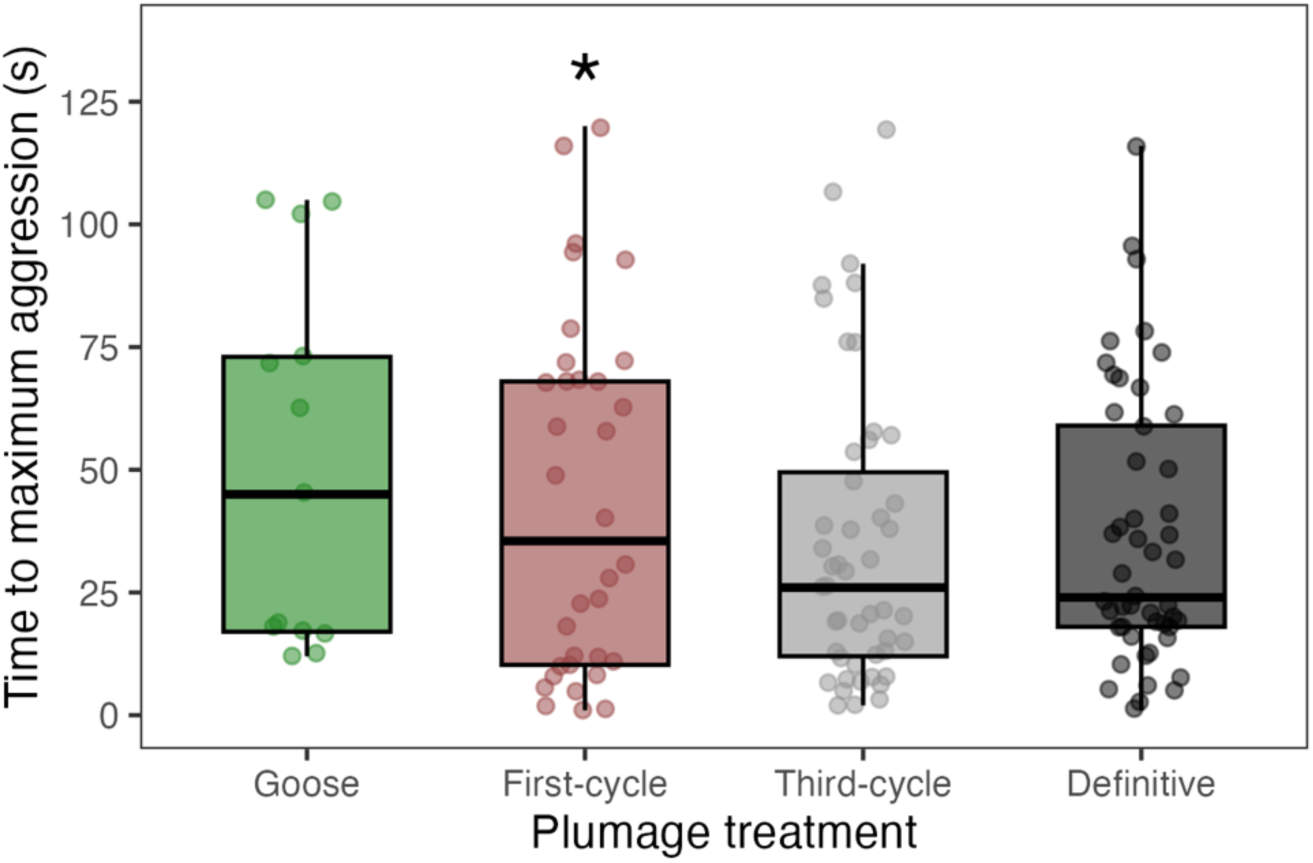
Aggressive American Herring Gulls responded more slowly toward first-cycle gull plumage compared to definitive gull plumage. Only trials with an aggressive response score of 2 or higher were included, limiting the sample size for the goose control. Asterisks indicate p <0.05 (*) compared to the intercept category of definitive plumage, from a linear mixed effects model.

**Table 4:** Results of linear model for time until maximum aggression (s). Definitive gull plumage treatment is the intercept category.

| **Term** | **Estimate** | **S.E.** | **DF** | ***t*** | ***p*** |
| --- | --- | --- | --- | --- | --- |
| (Intercept) | 49.22 | 9.44 | 85 | 5.22 | 0.00 |
| Plumage – Third-cycle | 0.92 | 6.03 | 49 | 0.15 | 0.88 |
| Plumage – First-cycle | 14.32 | 6.79 | 49 | 2.11 | **0.04** |
| Plumage – Goose | 18.28 | 9.87 | 49 | 1.85 | 0.07 |
| Trial order – 2 | -10.29 | 7.32 | 49 | -1.41 | 0.17 |
| Trial order – 3 | -16.23 | 7.20 | 49 | -2.26 | **0.03** |
| Trial order – 4 | -20.86 | 7.03 | 49 | -2.97 | **0.01** |
| Num. parents – 2 | 18.15 | 6.73 | 49 | 2.70 | **0.01** |
| Max. aggression – 3 | -3.50 | 8.37 | 49 | -0.42 | 0.68 |
| Max. aggression – 4 | -9.73 | 8.09 | 49 | -1.20 | 0.23 |

Aside from plumage, gulls reached peak aggression most slowly in the first trial (47.4 ± 35.2 s), slightly faster in the second (42.8 ± 30.5 s), and much faster at the third (33.4 ± 31.4 s) and fourth trials (34.3 ± 29.1 s; Table 4; Fig. S5), with the increases for third and fourth trials being statistically significant. Gulls also responded more slowly when two parents were present (53.2 ± 34.6 s) compared to one parent (36.1 ± 30.2 s; Table 4). Response times did not differ based on level of aggression (Table 4).

## Discussion

Multiple orders of seabirds have evolved prolonged delayed plumage maturation (Hawkins et al. 2012), but few studies have investigated the evolutionary significance of predefinitive plumages in these species. Breeding American Herring Gulls responded toward first-cycle predefinitive plumage stimuli with less frequent aggression (Fig. 2; Table 2), lower aggression levels (Fig. 3; Table 3), and slower aggressive reaction times (Fig. 4; Table 4) compared to definitive adult plumage stimuli. Our experiment with American Herring Gulls supports the hypothesis that substantial brown plumage elements, resembling those of first-cycle birds (∼12 months old), act as social signals that can reduce the risk of conflict with older, breeding birds (Taylor 2024).

These results are consistent with previous studies of delayed plumage maturation that find predefinitive plumages decrease aggression toward young birds, such as for lekking manakins and territorial songbirds (Lyon and Montgomerie 1986; McDonald 1993; Hawkins et al. 2012; Schaedler et al. 2021). These studies have focused on sexually dimorphic species, where female predefinitive plumages generally reduce competition among young males that are beginning to compete for territories or display opportunities. In contrast, delayed plumage maturation in seabirds is sexually monomorphic, with predefinitive plumages that are unique to immature birds rather than resembling a particular sex. This pattern suggests that delayed plumage maturation in seabirds has evolved in line with their social systems, which typically involve social monogamy with biparental territoriality and offspring care (Cockburn 2006). A promising opportunity for future studies of delayed development would involve comparing the dynamics of delayed plumage maturation across sexually dimorphic lineages (e.g., some songbirds or lekking birds) versus sexually monomorphic lineages (e.g., some seabirds or large raptors).

In many gull species, predefinitive plumages have mottled brown and black elements that resemble the natal and juvenile plumages of chicks and recent fledglings. Thus, our results indicate that immature gulls have co-opted signals from chick plumages, which may aid in crypsis (Tinbergen 1961, Hill and McGraw 2006) or may possibly help to reduce aggression toward vulnerable chicks themselves (Pierotti and Murphy 1987, Nelson 1978). A plumage that reduces the risk of aggression may help young birds navigate the social environment of the breeding colony and allow young animals to develop the breeding and territorial skills necessary to begin breeding themselves (Taylor 2024). Young birds of many seabird species return to colonies for years before breeding (Pickering 1966, Nelson 1978), despite colonies being dense social environments with frequent competition and physical aggression (Coulson 2002). Delayed plumage maturation, and the reduction of aggression toward young gulls, may allow young birds to engage with the social contexts without suffering injury.

Notably, our experiments found no difference in aggression toward the third-cycle plumage treatment compared to the definitive plumage treatment. Third-cycle gulls are the most common predefinitive birds in American Herring Gull breeding colonies, whereas first-cycle and second-cycle birds are less frequent (Taylor 2024). There are at least three potential explanations for this disparity. First, third-cycle birds may genuinely lack the plumage signals that reduce aggression in younger predefinitive birds, instead developing enough definitive traits that their plumages no longer function as signals of immaturity. For example, third-cycle birds exhibit a red gonydeal spot, which is used in adult signaling to chicks (Fig. 1A vs. B; Tinbergen and Perdeck 1950; Tinbergen 1953). From this perspective, predefinitive plumages may not serve as social signals for the third-cycle birds that return to colonies, but rather function as signals for younger birds that do not return to colonies and instead navigate other social environments such as foraging flocks or non-breeding grounds (Ulfstrand 1979, Wakefield et al. 2019). A second possibility is that our approach to quantifying aggression, using five broad categories, obscured key differences in reaction to the third-cycle plumage treatment. Our methods focused on obvious signals of aggression but did not consider postural cues (such as extending or contracting the neck) that play important roles in gull social interactions (Tinbergen 1961; Stout and Brass 1969, MacLean and Bonter 2013). Thus, our results may overlook more subtle variations in the response of breeding gulls towards all our visual stimuli. Finally, a third possibility is that our artificial stimuli may have failed to represent salient features of third-cycle plumages. For example, much of the brown plumage retained on third-cycle gulls is visible only on the spread wing (Fig. S6) and was consequently not visible on our folded-wing models. Thus, gulls may manipulate the signaling properties of predefinitive plumage through opening their wings or other postural cues.

The relationships between physical appearance, posture, and behavior are difficult to disentangle. Indeed, these relationships have confounded our understanding of delayed plumage maturation in seabirds. Previous observers noted that young gulls with brown plumages are attacked more frequently by breeding adults, not less (Huntington 1954; Ulfstrand 1979; Taylor 2024). However, they were unable to distinguish the effects of plumage and behavior using observational methods alone. Following classic work on visual signaling (Tinbergen and Perdeck 1950, Jones 1990), posture (Stout and Brass 1969, Galusha and Stout 1977), and nest defense (Shah et al. 2025) our study suggests that visual stimulus experiments in gulls can offer new insights into how social aggression evolves in the complicated context of a breeding colony. Our experimental investigation supports the idea that predefinitive plumages reduce aggression in the complex, competitive social context of a breeding colony.

## Acknowledgements

We are indebted to Patricia Jones, Ian Kyle, and the Bowdoin Scientific Station on Kent Island for providing access to the Kent Island breeding colony. We thank Tracey and Nicole Faber for painting the models. We are grateful to Richard Prum for discussions on the topic. Kristof Zyskowski provided access to reference specimens at the Yale Peabody Museum of Natural History. This study was funded by the Yale Institute for Biospheric Studies and the Yale School of the Environment.

## Notes

### Competing Interest Statement

The authors have declared no competing interest.

### Summary of Updates

The acknowledgements section, which was missing before, has been added, and the copyright was changed to CC-BY-NC.

## References

1. Anderson, C. M., Gilchrist, H. G., Ronconi, R. A., Shlepr, K. R., Clark, D. E., Weseloh, D. V. C., Robertson, G. J., & Mallory, M. L. (2019). Winter home range and habitat selection differs among breeding populations of herring gulls in eastern North America. Movement Ecology, 7, 8. 10.1186/s40462-019-0152-x.

2. Bates, D., Mächler, M., Bolker, B., & Walker, S. (2015). Fitting linear mixed-effects models using lme4. Journal of Statistical Software, 67(1), 1–48. 10.18637/jss.v067.i01.

3. Bennett, J., Jamieson, E., Ronconi, R., & Wong, S. (2017). Variability in egg size and population declines of Herring Gulls in relation to fisheries and climate conditions. Avian Conservation and Ecology, 12(2), 16. 10.5751/ACE-01118-120216.

4. Chabrzyk, G. & Coulson, J. C. (1976). Survival and recruitment in the herring gull *Larus argentatus*. Journal of Animal Ecology, 45(1), 187–203.

5. Christensen, R (2023). Ordinal-regression models for ordinal data. R package version 2023.12-4.1, <https://CRAN.R-project.org/package=ordinal>.

6. Cockburn, A. (2006). Prevalence of different models of parental care in birds. Proceedings of the Royal Society B, 273, 1375–1383. 10.1098/rspb.2005.3458.

7. Coulson, J. C. (2002). Colonial breeding in seabirds. Biology of marine birds, 87–113.

8. Galusha, J. G. & Stout, J. F. (1977). Aggressive communication by *Larus glaucescens* Part IV: Experiments on visual communication. Behavior, 62(3), 222–235.

9. Grant, P. J. (1982). Gulls: A guide to identification. Academic Press.

10. Gross, A. O. (1940). The migration of Kent Island herring gulls. Bird-Banding, 11(4), 129–155. 10.2307/4509634.

11. Hawkins, G. L., Hill, G. E., & Mercadante, A. (2012). Delayed plumage maturation and delayed reproductive investment in birds. Biological Reviews, 87(2), 257–274.

12. Hill, G. E., & McGraw, K. J. (Eds.) (2006). Bird coloration (vol. 2; pp. 201–232). Harvard University Press.

13. Huntington, C. (1954). Age discrimination in a breeding colony of the herring gull Larus argentatus. Acta XI Congressus Internationalis Ornithologici, 467-469.

14. Jones, I. L. (1990). Plumage variability functions for status signalling in least auklets. Animal Behaviour, 39, 967–975.

15. Kralick, A. E., O’Connell, C. A., Bastian, M. L., Hoke, M. K., Zemel, B. S., Schurr, T. G., & Tocheri, M. W. (2023). Beyond dimorphism: Body size variation among adult orangutans is not dichotomous by sex. Integrative and Comparative Biology, 63(4), 907–921.

16. Lyon, B. E. & Montgomerie, R. D. (1986). Delayed plumage maturation in passerine birds: Reliable signaling by subordinate males? Evolution, 40(3), 605–615.

17. MacLean, S. A. & Bonter, D. N. (2013). The sound of danger: Threat sensitivity to predator vocalizations, alarm calls, and novelty in gulls. PLOS One, 8(12), e82384.

18. McDonald, D. B. (1993). Delayed plumage maturation and orderly queues for status: A manakin mannequin experiment. Ethology, 94(1): 31–45.

19. Muehter, V. R., Greene, E., & Ratcliffe, L. (1997). Delayed plumage maturation in Lazuli buntings: tests of the female mimicry and status signalling hypotheses. Behavioral Ecology and Sociobiology, 41, 281–290.

20. Nelson, J. B. (1966). The breeding biology of the gannet *Sula bassana* on the Bass Rock, Scotland. Ibis, 108, 584–626.

21. Nelson, J. B. (1978). The Gannet. T. & A. D. Poyser Limited. p.24-25.

22. Nisbet, I. C. T., Weseloh, D. V., Hebert, C. E., Mallory, M. L., Poole, A. F., Ellis, J. C., Pyle, P. & Patten, M. A. (2017). Herring gull (Larus argentatus), version 3.0. In *The Birds of North America* (P. G. Rodewald, Ed.). Cornell Lab of Ornithology, Ithaca, NY, USA. 10.2173/bna.hergul.03

23. Pickering, S. P. C. (1989). Attendance patterns and behaviour in relation to experience and pair-bond formation in the Wandering Albatross *Diomedea exulans* at South Georgia. Ibis, 131(2), 183–195. https://onlinelibrary.wiley.com/doi/abs/10.1111/j.1474-919X.1989.tb02761.x

24. Pierotti, R. & Annett, C. (1994). Patterns of aggression in gulls: asymmetris and tactics in different social categories. The Condor: Ornithological Applications, 96(3), 590–599. 10.2307/1369461.

25. Pierotti, R. & Murphy, E. C. (1987). Intergenerational conflicts in gulls. Animal Behaviour, 35(2), 435–444. 10.1016/S0003-3472(87)80268-3.

26. Pinheiro, J. & Bates, D., R Core Team (2024). _nlme: Linear and Nonlinear Mixed Effects Models_. R package version 3.1–166, <https://CRAN.R-project.org/package=nlme>.

27. Schaedler, L. M., Taylor, L. U., Prum, R. O., & Aciães, M. (2021). Constraint and function in the predefinitive plumages of manakins (Aves: Pipridae). Integrative and Comparative Biology, 61(4), 1363–1377.

28. Schreiber, R.W. (1970). Breeding Biology of Western Gulls (Larus occidentalis) on San Nicolas Island, California, 1968. The Condor, 72(2), 133–140.

29. Shah, S. S., Ellms, M. E., Dygean, F., Gunasekera, S., Hart, E., Ivanyi, R. E., Lane, M., Liu, D., Pitcher, D., Saucier, A., Schroeder, K. F., Triquet, J., Williams, K. M., Covino, K. M. (2025). Investigating diurnal effects and joint nest defense behavior in Herring Gulls. bioRxiv. 10.1101/2025.06.13.659635.

30. Stout, J. F., and Brass, M. E. (1969). Aggressive Communication by *Larus glaucescens:* Part II. Visual Communication. Behavior, 34(1/2), 42–54.

31. Taylor, L. U. (2024). Young American Herring Gulls (*Larus argentatus* subsp. Smithsonianus) have the opportunity for social development at the breeding colony. Waterbirds, 47(2): 1–12.

32. Tickell, W. L. N. (2000). Albatrosses. Yale University Press.

33. Tinbergen, N. (1953). The herring gull’s world: a study of the social behavior of birds. Frederick A. Praeger, Inc.

34. Tinbergen, N. (1961). The Herring Gull’s world: Study of the social behavior of birds. Basic Books.

35. Tinbergen, N., & Perdeck, A. C. (1950). On the stimulus situation releasing the begging response in the newly hatched Herring Gull chick (*Larus Argentatus Argentatus Pont*.). Behavior, 3(1), 1–39.

36. Ulfstrand, S. (1979). Age and plumage associated differences of behaviour among blackheaded gulls *Larus ridibundus*: foraging success, conflict victoriousness and reaction to disturbance. Oikos, 33(2), 160–166.

37. Wakefield, E. D., Furness, R. W., Lane, J. V., Jeglinski, J. W. E., & Pinder, S. J. (2019). Immature gannets follow adults in commuting flocks providing a potential mechanism for social learning. Journal of Avian Biology, 50(10). 10.1111/jav.02164.

38. Warner, R. R., & Swearer, S. E. (1991). Social control of sex change in the bluehead wrase, *Thalassoma bifasciatum* (Pisces: Labridae). Biol. Bull. 181, 199–204.

39. Wickham, H., Averick, M., Bryan, J., Chang, W., McGowan, L.D., François, R., Grolemnd, G., Hayes, A., Henry, L., Hester, J., Kuhn, M., Pedersen, T.L., Miller, E., Bache, S.M., Müller, K., Ooms, J., Robinson, D., Aeidel, D.P., Spinu, V., Takahashi, K., Vaughan, D., Wilke, C., Woo, K., & Yutani, H. (2019). Welcome to the Tidyverse. Journal of Open Source Software, 4(43), 1686.

