## Supplementary Data for "Experimental Study of Social Signaling through Delayed Plumage Maturation in a Colony-Nesting Seabird"

**for**

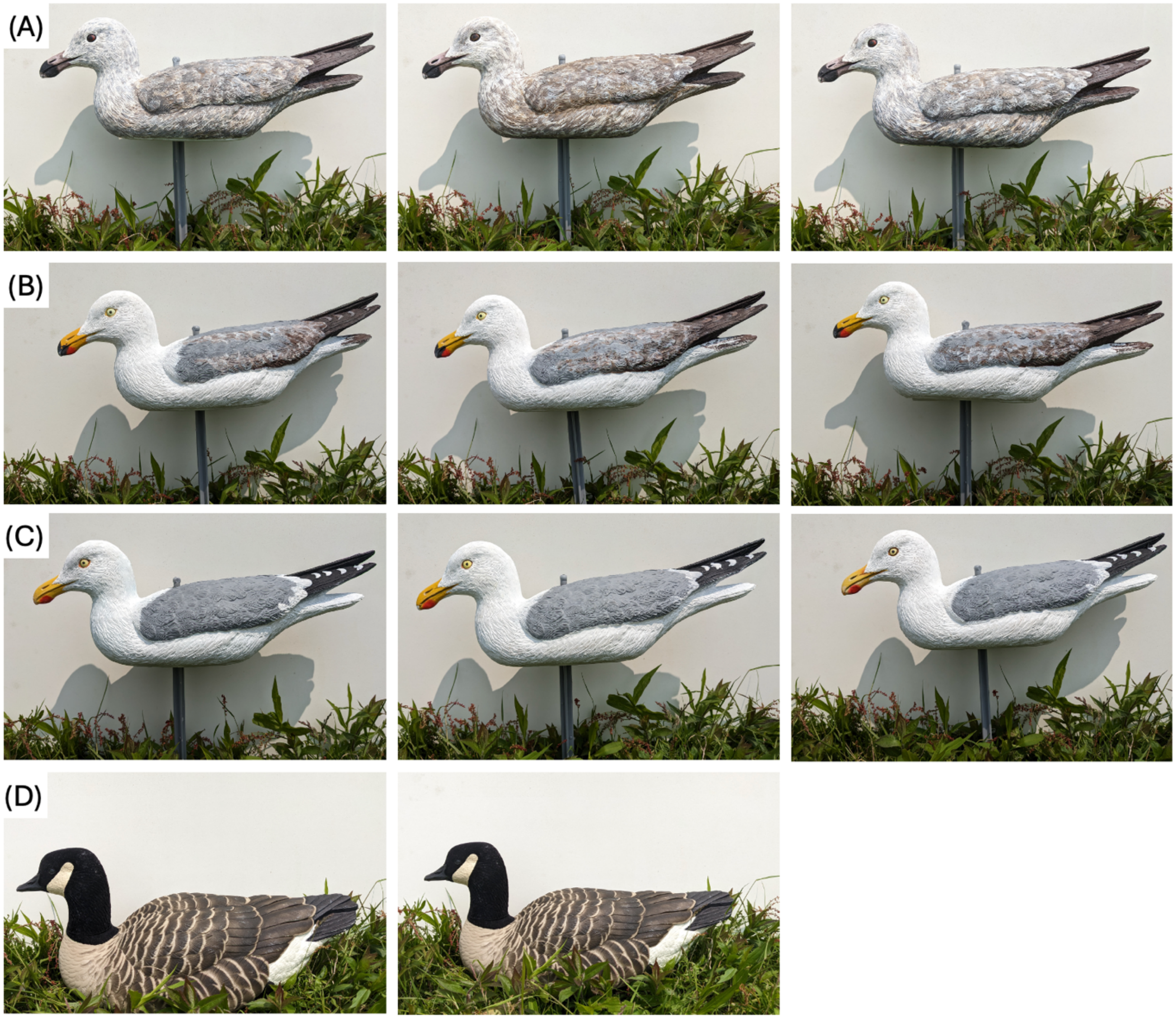


**Figure S1.** Exemplars of all plumage treatments. (A) Definitive gull plumage models. (B) Third-cycle gull plumage models. (C) First-cycle gull plumage models. (D) Goose control models.

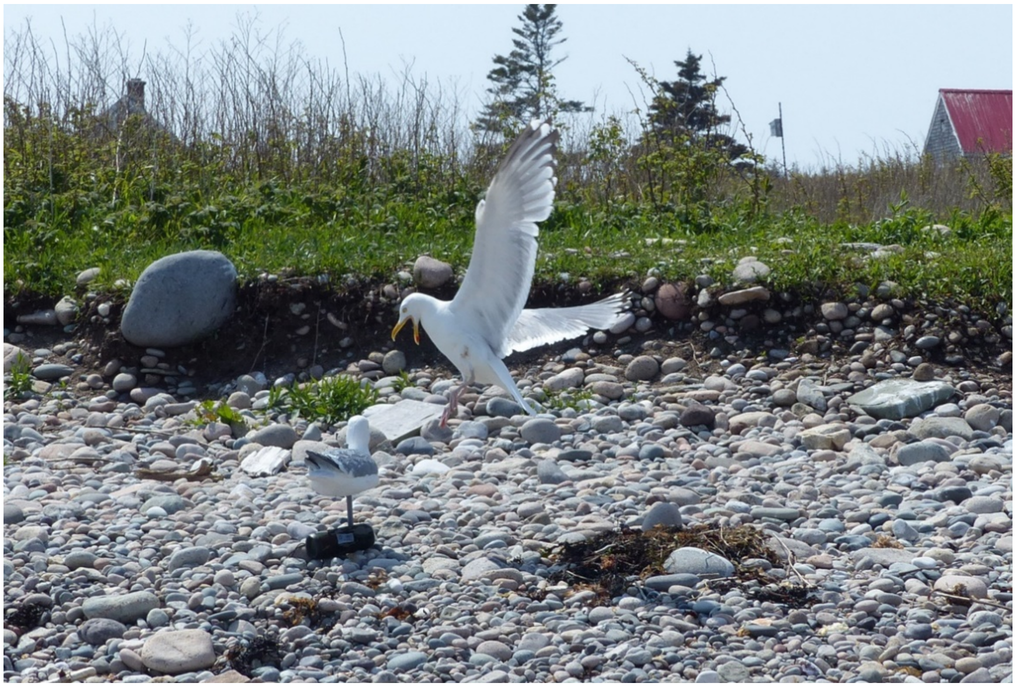

**Figure S2.** Example photograph of the experimental design on Kent Island. The visual stimulus (here, definitive gull plumage) and secondary audio stimulus (playback speaker) are placed 0.75 m to the left of the nest.

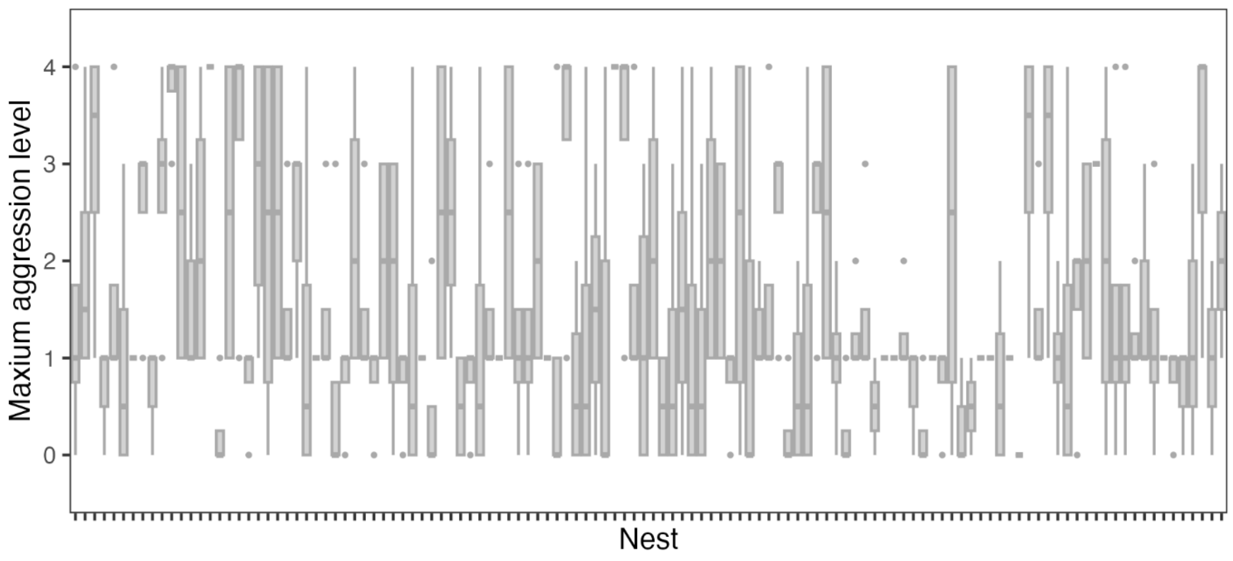


**Figure S3**. Variation in aggression across individual nests (the random effect in all models). There were four trials at each of 120 unique nests (total 480 trials, with 442 trials retained for analysis). Nests are ordered by day of year for the first trial at each nest, across both 2023 and 2024.

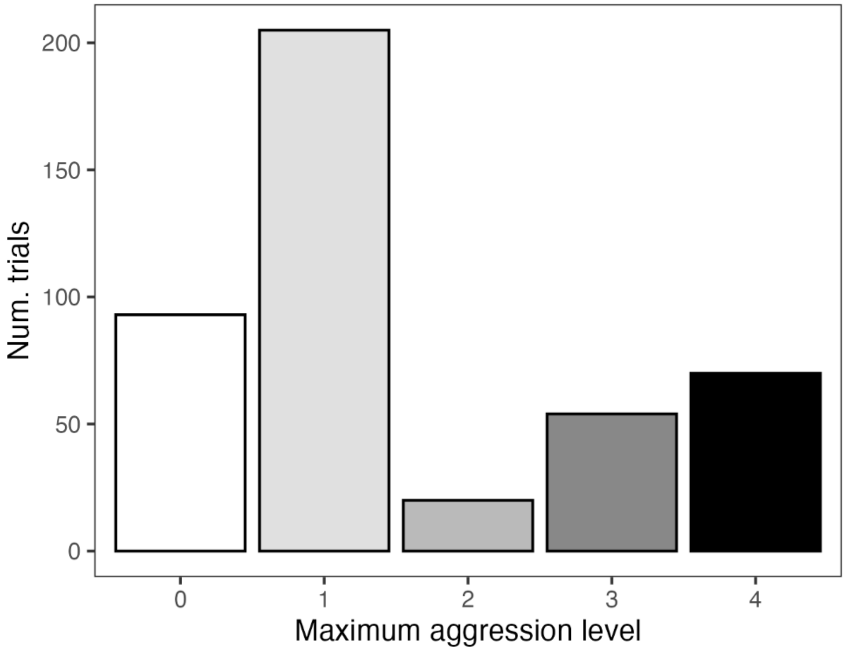


**Figure S4.** Histogram of number of trials that reached each of the five aggression categories.

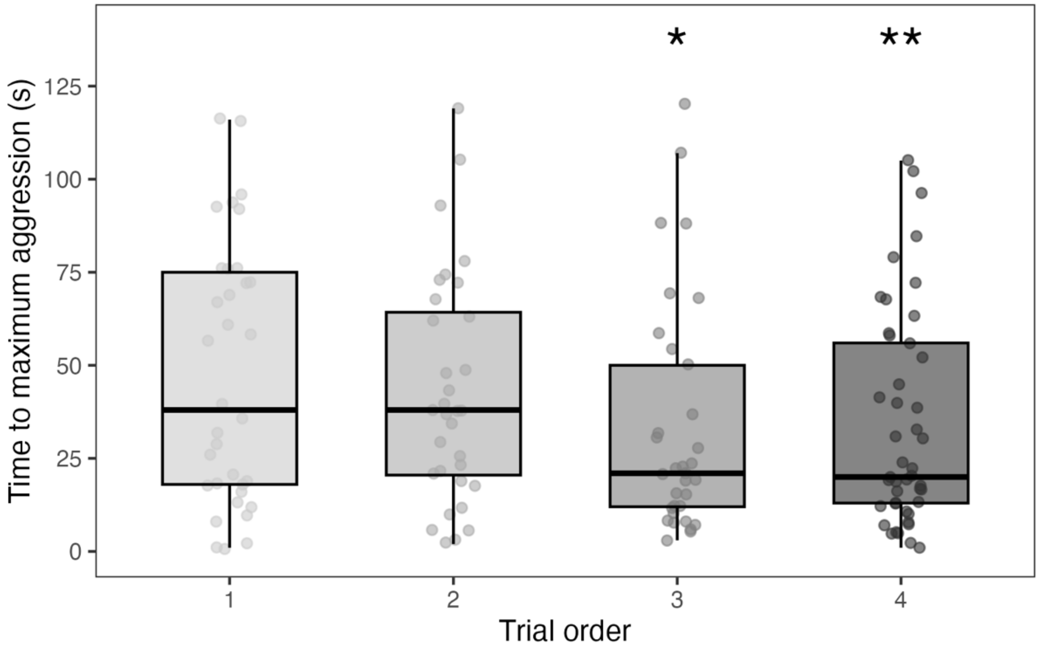


**Figure S5.** For trials with an aggressive response, gulls reacted significantly faster during third and fourth trials compared to the first trials. Asterisks indicate significance compared to the intercept category of trial order 1, from a linear mixed effects model (* $p < 0.05$, ** < 0.01).

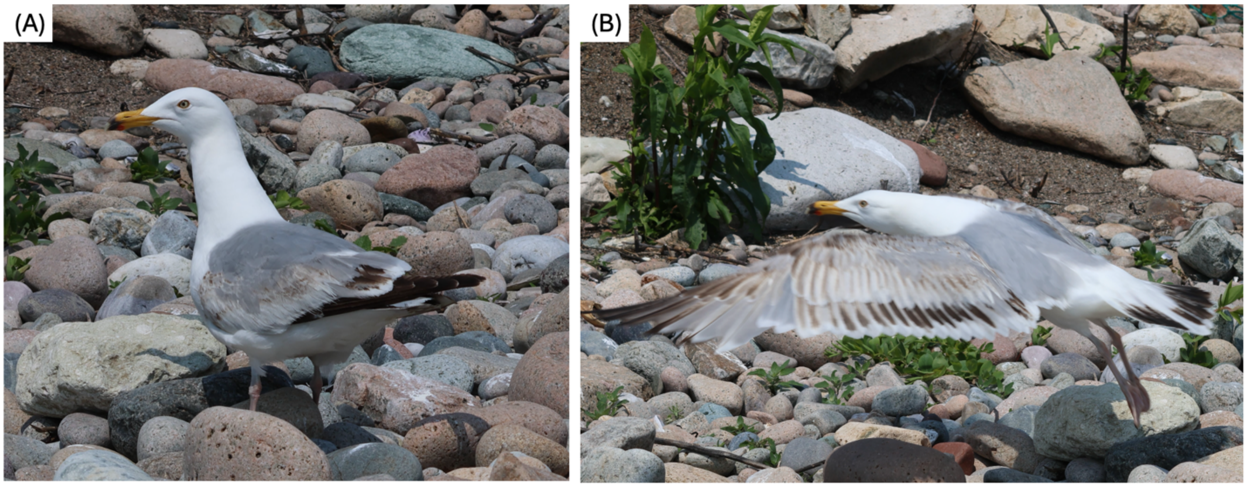


**Figure S6.** Role of posture in predefinitive plumage displays. (A) A third-cycle gull photographed on Kent Island, with limited brown visible on the folded wings. (B) The same bird photographed with spread wings, displaying much more mottled brown plumage on the wings as well as more black in the tail.
